# Exogene: A performant workflow for detecting viral integrations from paired-end next-generation sequencing data

**DOI:** 10.1101/2021.04.19.440427

**Authors:** Zachary Stephens, Daniel O’Brien, Mrunal Dehankar, Lewis R. Roberts, Ravishankar K. Iyer, Jean-Pierre Kocher

**Affiliations:** Department of Electrical and Computer Engineering, University of Illinois Urbana-Champaign, Urbana, IL, USA; Department of Health Sciences Research, Mayo Clinic, Rochester, MN, USA; Department of Internal Medicine, Mayo Clinic, Rochester, MN, USA

## Abstract

The integration of viruses into the human genome is known to be associated with tumorigenesis in many cancers, but the accurate detection of integration breakpoints from short read sequencing data is made difficult by human-viral homologies, viral genome heterogeneity, coverage limitations, and other factors. To address this, we present Exogene, a sensitive and efficient workflow for detecting viral integrations from paired-end next generation sequencing data. Exogene’s read filtering and breakpoint detection strategies yield integration coordinates that are highly concordant with those found in long read validation sets. We demonstrate this concordance across 6 TCGA Hepatocellular carcinoma (HCC) tumor samples, identifying integrations of hepatitis B virus that are validated by long reads. Additionally, we applied Exogene to targeted capture data from 426 previously studied HCC samples, achieving 98.9% concordance with existing methods and identifying 238 high-confidence integrations that were not previously reported. Exogene is applicable to multiple types of paired-end sequence data, including genome, exome, RNA-Seq or targeted capture.

## Introduction

The integration of viruses into the human genome has been extensively studied and is central to the etiology of many prominent diseases [1, 2]. The link between viral integration and tumorigenesis in humans was established in the 1960s [3, 4], and since then there has been increasing experimental evidence associating such integrations with human cancers. Examples include human papilloma virus and cervical cancers [5], hepatitis B viruses (HBV) and liver cancer [6], herpes and Epstein-Barr viruses and lymphoma [7, 8], among others [9–11]. Over the last decade, next generation sequencing (NGS) technologies have accelerated the study of viral integration, enhancing our understanding of virus-associated tumor development and enabling the study of viral integration at genome-wide scales. These studies have found many associations between viral integration and host genome instability, e.g. regions surrounding integration sites exhibiting increased mutation rates, copy number alterations, or aberrant gene expression [12–14]. Additionally, it has been observed that viral integrations in tumor samples are often enriched near genes with known associations to cancer, including *MLL4* [15], *MYC* [16] and *TERT* [17].

Despite their clinical utility, the sensitive detection of human/viral junctions from NGS data is made challenging by several factors. These include sequence similarities between host and viral genomes, integrated virus segments that differ from available reference genomes, and the limited number of validated integration sites in publicly available samples that can be used to assess detection accuracies. Several software applications for detecting viral integrations have been recently reviewed [18, 19], with each tool generally starting from unmapped reads or from reads mapped to a combined human + viral reference database. The tuning of read filtering and breakpoint detection strategies is crucial for the efficient extraction of informative reads, particularly when working with tumor samples where the number of reads supporting an integration may be limited. Additionally, these methods must be computationally efficient to be useful in practice, and must be scalable to the size and complexity of large sequencing datasets.

To address these challenges, we present Exogene, a new workflow for reporting viral integration sites from paired-end sequencing data. Exogene is computationally efficient and can identify integration coordinates from paired-end whole-genome (WGS), whole-exome (WES), RNA-Seq, or targeted capture data. Exogene builds upon our previous methodology HGT-ID [20], with new preprocessing, alignment, and filtering strategies to improve breakpoint precision.

We demonstrate Exogene’s ability to identify viral integration sites in 6 samples (5 WES, and 1 WGS + WES) from the TCGA Liver Hepatocellular Carcinoma (HCC) project. We show that the coordinates reported by Exogene are highly concordant with those found in a long read validation set. We demonstrate an improvement in accuracy over HGT-ID, attributable to Exogene’s improved extraction of informative read pairs. Additionally, we demonstrate Exogene’s applicability to targeted capture data by processing 426 HCC tumor/normal pairs from a previous study, achieving 98.9% concordance with existing results and augmenting them with 238 novel high-confidence integrations. Exogene’s runtime scales with input file size, and can process a 100× coverage WGS BAM (∼470 GB) within 12 hours (4 CPUs, 32GB memory).

Exogene is distributed as a Docker container, and is available at github.com/zstephens/exogene.

## Materials and methods

Exogene takes as input a BAM file, or alternately paired FASTQ files, and produces an output report of all detected integrations, including breakpoints, quality metrics and visualizations (Fig 1).

**Fig 1.**
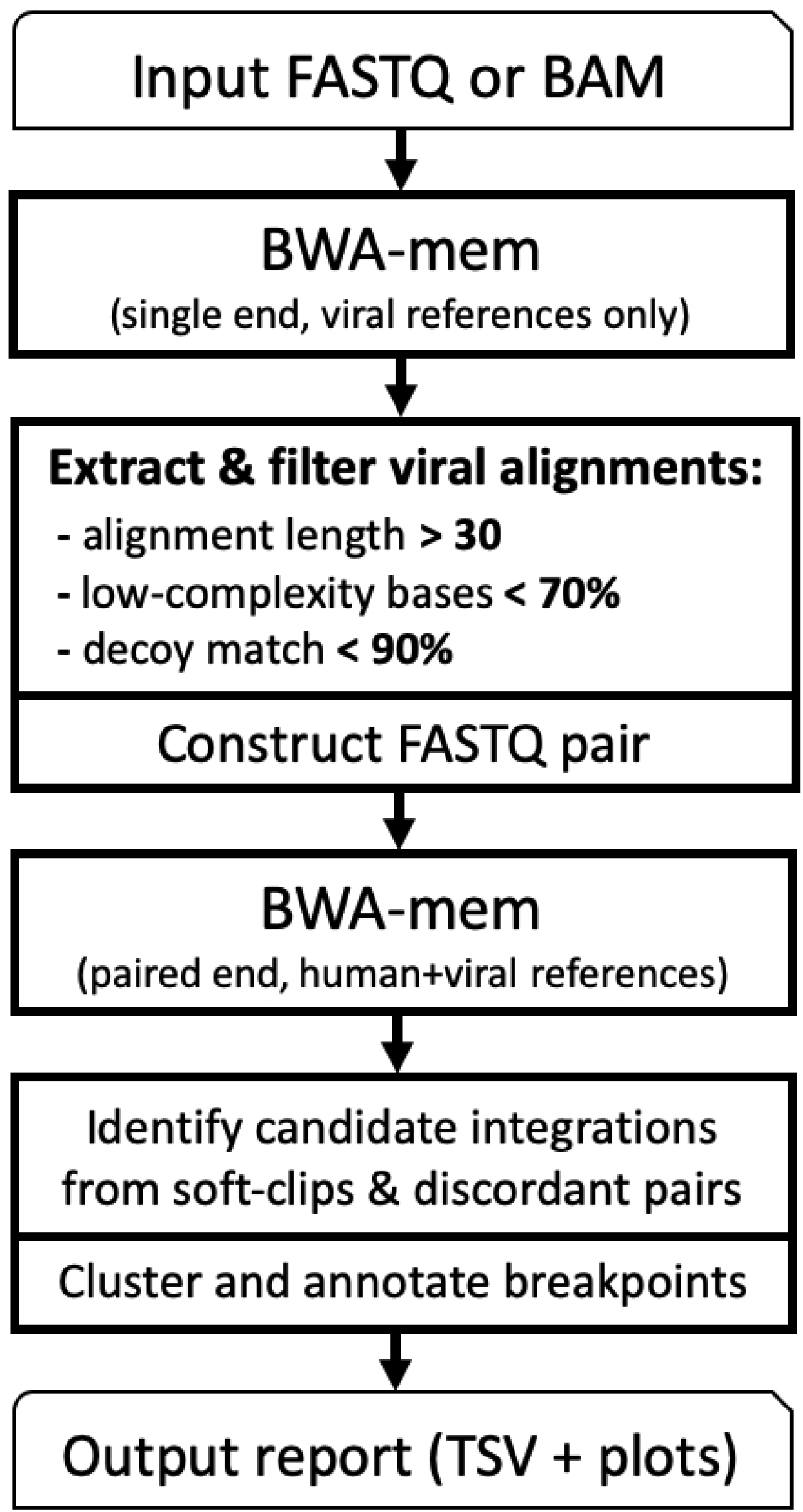
Overview of Exogene workflow

Exogene begins by aligning the input reads to a collection of 1,628 viral reference sequences that are included with the workflow. This is performed using BWA MEM in single-end mode. From the resulting BAM file we enumerate the names of all reads which were able to be mapped to a virus with an alignment length of at least *K*. By default Exogene uses *K* = 30, but for shorter reads it may be necessary to reduce this value. Because this first step maps all input reads to solely viral references, it will likely contain alignments of human DNA which were only mapped to a viral reference due to human/viral sequence similarity. We find that a vast majority of these reads are either low-complexity, or originate from regions of the human genome which we have apriori identified as having similar sequence content to one of the viruses. To address this, aligned reads are annotated for low-complexity sequence using Dustmasker [21]: If > *D*% of a read’s length is masked then it is discarded. By default Exogene uses *D* = 70%. Next, the reads are tested for similarity to a collection of decoy and transcriptome sequences (including exon-exon junctions), and reads are discarded if *> T* % of their length match in a single alignment. By default Exogene uses *T* = 90%.

Two FASTQ files are then constructed by extracting, from the original BAM/ FASTQ, all read pairs in which one or more mates are aligned to a virus and passed all filters. These reads are then aligned using BWA MEM in paired-end mode to the combined human+viral reference (human reference build GRCh38). BWA is run with the -Y input option so that large soft-clipped segments are recorded as supplementary alignments.

SAM records are extracted from this alignment if their associated read pair has at least one alignment to human and at least one alignment to virus. Possible integration coordinates are identified from soft-clipping in human alignments, and the specific virus is inferred from viral alignments (either from a supplementary alignment of the read containing the soft-clip, or from the primary alignment of its mate). If no clipping is present then we only have the discordant mapping as evidence of integration. In this case integration coordinates are estimated based on the position of the human alignment and fragment length statistics provided by BWA. In the event that one or more of the reads is multi-mapped, that is, aligned at multiple positions with mapping quality 0, a “representative” alignment is chosen for each read (see Supplementary Fig S1).

Detected integrations are clustered by position and each cluster is summarized with its predicted integration coordinate, supporting read count, and quality metrics such as breakpoint variance and read mapping quality (MAPQ) distribution. If desired, the user can specify to include weakly-supported integrations in the final output report, which include integrations flagged as:

- **Low read count**: less than *N*_*s*_ soft-clipped reads, less than *N*_*d*_ discordant reads. By default *N*_*s*_ = 2, *N*_*d*_ = 5.
- **Low MAPQ**: Supporting reads were aligned with mapping quality 0. This filter only applies to reads mapped to human. It is expected that viral alignments may have low mapping quality because our viral database contains many highly similar sequences for certain viruses.
- **Uncertain coordinate**: position is in large repetitive region, or in regions with high sequence similarity to viral references.

### Viral References

Exogene uses a database of 1,628 viral reference sequences. A majority of the sequences were sourced from Virus-Host DB ^1^, which compiles sequences from RefSeq, GenBank, EBI, UniProt, ViralZone, and published literature. We augmented the set with specific genomes of interest sourced from specialized databases. Most notably, additional strains of herpes and HPV (sourced from GenBank), and additional strains of HBV (genotypes A-H and various recombinants sourced from HBVdb [22]). We include multiple strains of certain viruses to increase the likelihood of extracting reads originating from viral genomes that may differ from the available reference sequences.

### Long Read Validation

To evaluate Exogene’s performance we compared its results to long reads sequenced from the same samples. DNA was extracted from frozen liver tumor tissue of 6 individuals from the TCGA Liver Hepatocellular Carcinoma project. Short reads were obtained from TCGA, including 1 WGS (barcode TCGA-DD-A1EL) and 6 WES (barcodes TCGA-DD-AACV, TCGA-DD-AAD0, TCGA-DD-AADL, TCGA-DD-AADU, TCGA-DD-AADV, and TCGA-DD-A1EL). The sequencing was performed at the Human Genome Sequencing Center (HGSC) at Baylor College of Medicine. Paired-end DNA sequence libraries were prepared following standard HGSC protocols ^2^.

Long reads were sequenced at Mayo Clinic on a PacBio Sequel II, following the standard protocols for Continuous Long Reads (CLR) and high-fidelity Circular Consensus Sequences (HiFi/CCS) reads ^3^. A 10kb fragment size was targeted for the HiFi reads, which were processed using the CCS application in SMRT Link v7.0 and requiring a minimum predicted accuracy of 99.9% per read.

Integration sites were identified in the long reads by aligning them to the combined human + viral references using pbmm2, a fork of the popular minimap2 aligner [23]. Reads with alignments to both human and viral sequence were extracted, and the position of the soft-clipped coordinates were used to validate Exogene’s reported integration sites.

## Results

We processed short reads from TCGA-DD-A1EL WGS through both Exogene and HGT-ID and enumerated all HBV integration sites that were also found in long reads (Table 1). On average, Exogene’s integration coordinates differed from long reads by 1.6 bp (std. 3.6 bp). HGT-ID differed by 175 pb (std. 102 bp). In addition to integration coordinates and quality metrics, Exogene produces figures showing the intersection of evidence at each integration site (example in Fig 2).

**Table 1.**
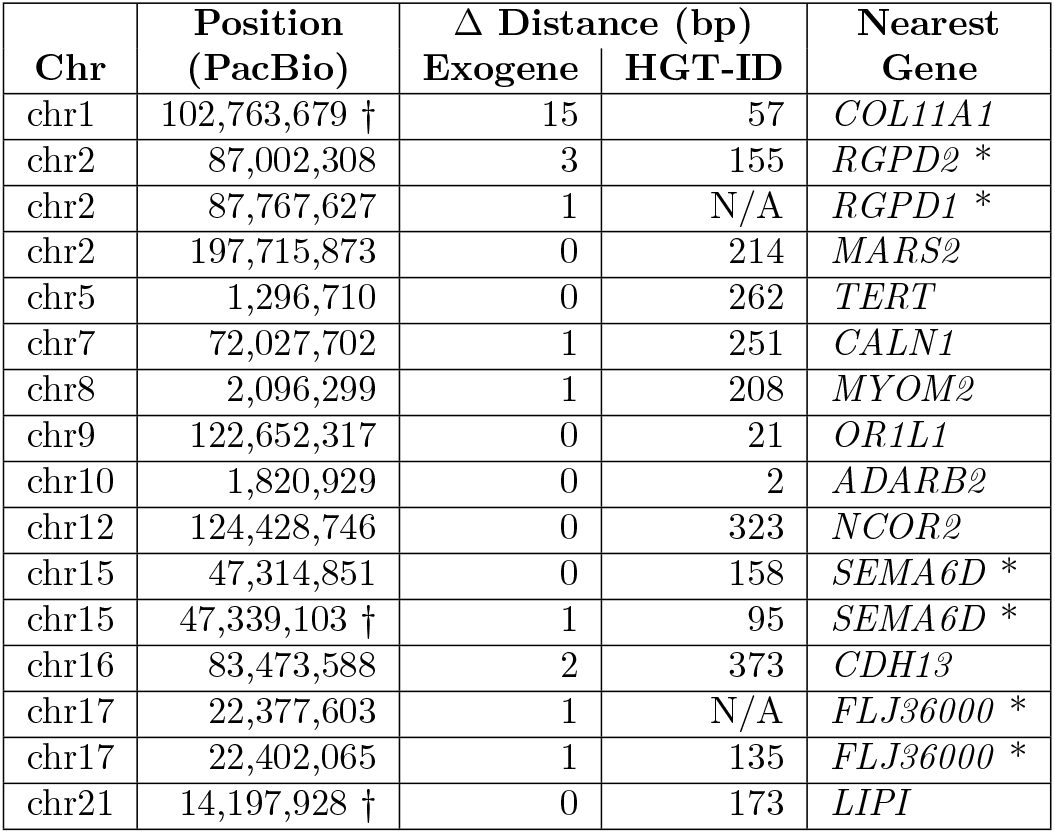
Overview of HBV integration sites in TCGA-DD-A1EL. Δ distance denotes the distance from integration sites reported by Exogene and HGT-ID to those found in long reads. “N/A” indicates an integration that was not reported by one of the short read workflows. * Breakpoints are in repeat regions and supporting reads have non-unique alignments. † Multiple integration coordinates were found within close proximity, in these cases Δ distance distance is computed as the distance from the short read coordinate to the nearest integration found in long reads.

**Fig 2.**
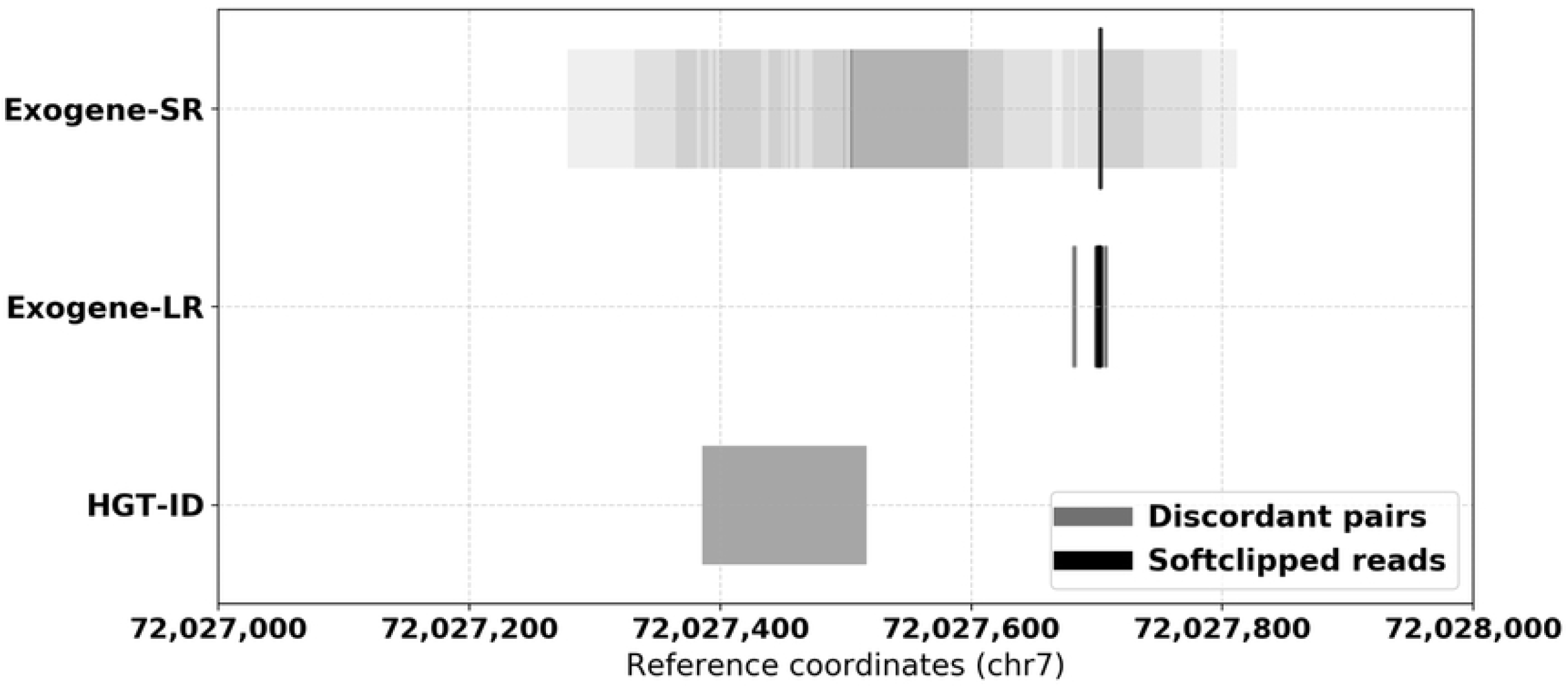
Comparison of evidence for HBV integration at chr7:72,027,703 from Exogene and HGT-ID. Shaded regions indicate breakpoint ranges as inferred from read fragment lengths and orientations, darker shades indicate greater support.

At 14/16 sites, both Exogene and HGT-ID reported an integration corroborated by long reads. At the remaining 2 sites, Exogene reported integrations that were missed by HGT-ID. We note that these 2 integrations were reported in repetitive regions of the genome near genes *RGPD1* and *FLJ360000*. The short reads that support these integrations were all aligned with mapping quality 0, indicating that they map equally well to other locations and thus the reported integration coordinate is likely not unique. The long reads, however, were aligned with high mapping quality, suggesting that the integrations are not false positives and that the size of the repetitive elements they are located in are larger than the length of the short reads, but smaller than the length of the long reads.

### Computational Performance

The A1EL WGS BAM was approximately 470 GB in size, which Exogene completed processing in 12 hours of runtime (4 threads, 48 CPU hrs in total). HGT-ID completed in 26 hours (4 threads, 41 CPU hours in total). Note that Exogene does not require an aligned BAM as input, so if we were starting with FASTQ files HGT-ID would require additional computational time to first align the reads. A majority of Exogene’s runtime is spent in the initial BWA alignment to viral references. Subsequent steps complete quickly as the subset of read pairs with viral alignments which pass all read filters is generally small as compared to the size of the original input BAM/FASTQ.

### Additional WES samples

Next we processed 6 WES samples with Exogene and identified 18 HBV integration sites with long read support (Table 2). HGT-ID was not included in this comparison as it only supports WGS and RNA-Seq input data. At 15/18 sites, Exogene reported integration coordinates within ≤ 2 bp of coordinates identified in long reads. Across all 18 sites, Exogene’s reported coordinates differed from long reads by 11.6 bp on average (std. 35.8 bp). Noteworthy integration sites include *TERT* promoter, which is well known to be associated with HCC. Integrations were also reported in *ADARB2, RALYL*, and *URI1*, which have been associated with liver tumor development [24–26].

**Table 2.** HBV integration sites in 6 WES samples. Δ distance is defined as the distance from integration sites reported by Exogene to those found in long reads. * Breakpoint in region with human/viral sequence similarity.

| ID | Chr | Integration Position | $\Delta$ Distance (bp) | Nearest Gene |
| --- | --- | --- | --- | --- |
| AACV | chr5 | 1,294,755 | 27 | *TERT* |
| AAD0 | chr5 | 1,295,109 | 7 | *TERT* |
| AADL | chr5 | 1,295,056 | 0 | *TERT* |
| AADL | chr8 | 8,338,904 | 157 | *PRAG1* * |
| AADU | chr2 | 29,900,177 | 0 | *ALK* |
| AADU | chr2 | 89,665,978 | 0 | *RPIA* * |
| AADU | chr5 | 111,687,499 | 2 | *STARD4-AS1* |
| AADU | chr8 | 83,576,876 | 0 | *RALYL* |
| AADV | chr17 | 43,747,355 | 1 | *SOST* |
| AADV | chr19 | 29,936,802 | 2 | *URI1* |
| A1EL | chr1 | 102,763,687 | 8 | *COL11A1* |
| A1EL | chr2 | 87,767,626 | 1 | *RGPD1* |
| A1EL | chr2 | 197,715,873 | 0 | *MARS2* |
| A1EL | chr8 | 2,096,300 | 1 | *MYOM2* |
| A1EL | chr10 | 1,820,929 | 0 | *ADARB2* |
| A1EL | chr15 | 47,314,851 | 0 | *SEMA6D* |
| A1EL | chr16 | 83,473,590 | 2 | *CDH13* |
| A1EL | chr17 | 22,402,064 | 1 | *FLJ36000* |

### Exogene Applied to Targeted Capture

To further validate Exogene, we apply it to short read targeted capture data sequenced for a previous study on HBV integrations in liver tumors [12]. For this study the authors designed sequence-capture probes for 8 strains of HBV, which they used to extract and sequence viral integration sites from liver tissue. The authors used the HIVID pipeline [27] to identify 4199 HBV integrations across 426 tumor/normal pairs. 707 of the 4199 integrations (16.8%) reported by HIVID were located in centromeres, telomeres, or other large repetitive regions of the genome where unique coordinates cannot be reliably inferred (i.e. regions where reads supporting a particular integration coordinate would align equally well to other positions in the reference genome). Thus we solely consider the 3492 integrations not reported in such regions.

We ran Exogene on each of the 426 tumor/normal pairs using paired-end FASTQ data hosted on the Sequence Read Archive [28] under project accession PRJNA298941. Exogene reported 3454/3492 (98.9%) of the integrations identified by HIVID. The full table of reported integrations is provided in Supplementary Fig S2. The average processing time for each sample was 20 minutes, and used 6 GB of memory.

Of these 3454 concordant calls, 3265 were supported by soft-clipped reads, and the remaining 189 had only discordant read pairs as evidence. Of the 3265 concordant calls with soft-clipped evidence, 2861 (87.6%) of the integration coordinates reported by Exogene were identical to those reported by HIVID. Integrations with non-identical coordinates between the two workflows differed by 48 bp on average. The coverage depth and mapping quality varied substantially in reads extracted by Exogene (Fig 3). That is, very few reads with high mapping quality were extracted at certain sites identified by HIVID as having an HBV integration. 1277/3454 (37.0%) concordant integrations had more than half of their supporting reads aligned with mapping quality 0, 780 of which were supported entirely by such reads. We note that these low-confidence integrations tend to occur in clusters, often near low-complexity regions. Genes most affected by this include *HERC2, CCDC144, SNORD3*, and *SLBP*.

**Fig 3.**
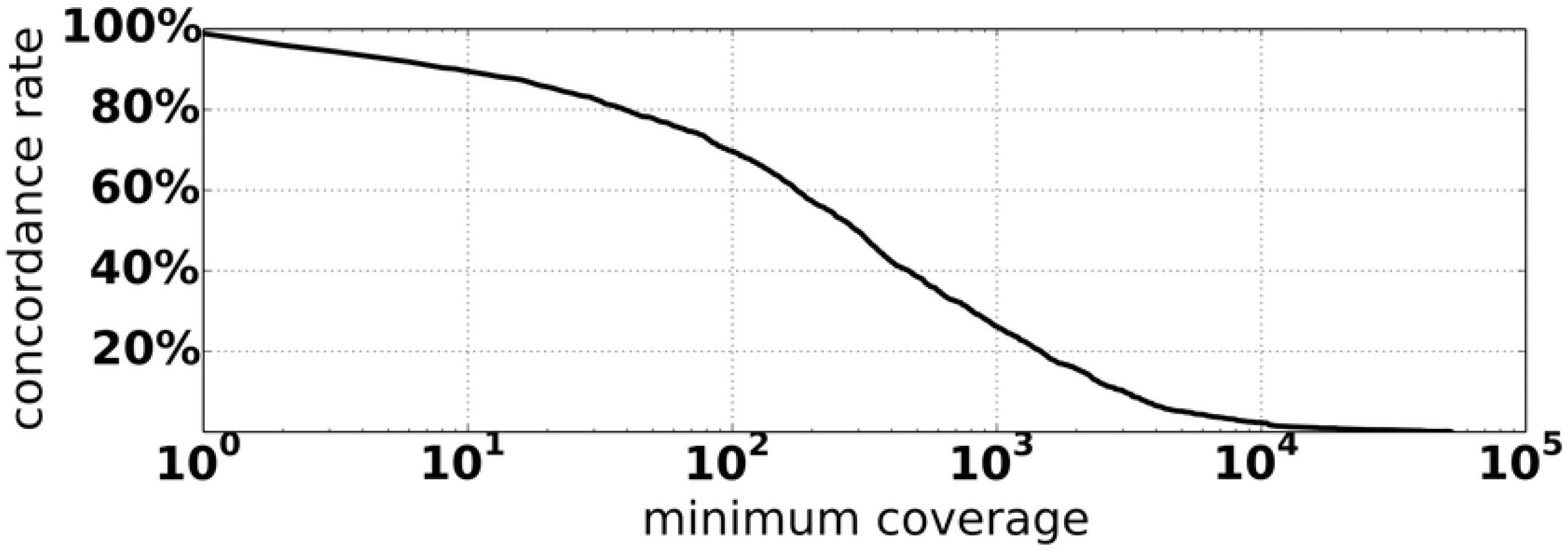
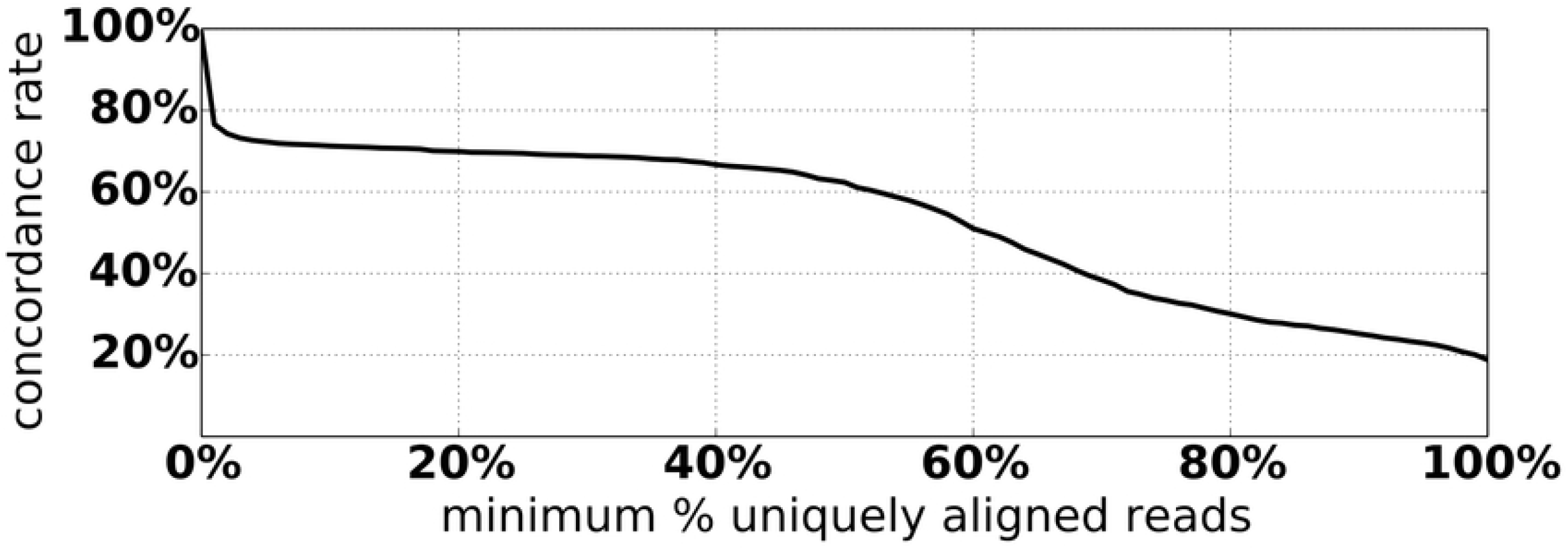
Concordance rate of Exogene and HIVID calls, as a function of minimum coverage and minimum allowable percentage of reads aligned with mapping quality 0.

Exogene reported additional HBV integrations that were not found in the HIVID results. Based on the distributions in Fig 3, we identified 238 novel integrations supported by at least 100 uniquely aligned reads. While these novel integrations are not enriched in any particular genomic region, a number of them hit introns of genes associated with HCC, including *WDHD1, THSD4*, and *KIF20A*.

From this comparison, we conclude that Exogene is effective on targeted capture data, achieving high concordance with the HIVID pipeline. Exogene’s annotations potentially reduce false positives in regions of poor mappability or human-viral sequence similarity by flagging integrations in these regions as low confidence. The novel integrations identified by Exogene are potentially valuable for future study.

## Discussion

Previously, many authors seeking to validate integration sites either compared against previous analyses of the same dataset [29, 30], or against PCR experiments on a limited number of sites [31]. Previous reviews have used simulated data to compare accuracy across methods [18, 19], but this approach is limited in its applicability to real samples which have additional complexities such as recombinant viral strains, confounding structural variation (including virus-mediated rearrangements), and sequencing biases that simulation tools do not replicate.

In addition to these strategies, another approach for validating integration sites is via intersecting results from multiple analyses on the same sample across different sequencing protocols or sequencing platforms. Long reads from ‘third-generation’ sequencers, such as those from PacBio or Oxford Nanopore, are attractive for this validation due to their increased ability to anchor large structural variation and to span repetitive genomic regions.

Using integrations identified from PacBio long reads as a baseline set, we compared results from Exogene to HGT-ID on one WGS sample with many integrations. We observe that on average, the breakpoints reported by Exogene-SR are significantly closer to those in long reads, as opposed to breakpoints reported by HGT-ID (Table 1). This is largely attributable to Exogene’s improved extraction of soft-clipped reads, which provide evidence for breakpoints at specific coordinates (as opposed to discordant read pairs, which support a range of possible breakpoint positions). Conversely, HGT-ID extracts most of its evidence from discordant read pairs and reports the average of their ranges as the final breakpoint. We attribute Exogene’s improved extraction of soft-clipped reads to three main factors: 1) The initial alignment to viral references only, instead of a combined human + viral FASTA. This ensures that reads of viral origin that would be preferentially aligned to human due to sequence similarity are retained for further analysis. 2) Instead of discarding reads with multiple alignments or alignments to blacklisted regions, we include them in reporting but flag them as low confidence. 3) Improved logic for choosing representative alignments in cases where reads are multi-mapped or have multiple supplementary alignments.

We observe similarly high concordance in the 6 WES samples, where at nearly every site the HBV integration coordinates reported by Exogene are very close to those found in long reads. There is only one site (near gene PRAG1) where the coordinates differ substantially. This is attributable to it being the only site where Exogene could not extract soft-clipped reads. When Exogene’s only source of evidence is discordant read pairs, the reported coordinate is estimated from alignment orientation and fragment length statistics (in a similar manner as HGT-ID).

### Usability

Workflows for identifying viral integrations typically leverage multiple third-party bioinformatics tools, sometimes requiring specific system configurations or laborious installation procedures. Additionally, it has been our experience that existing workflows exhibit poor stability, or that resource requirements make running them prohibitive. This has been commented on by other authors, who have excluded comparisons with certain tools due to an inability to successfully apply them to their samples [20, 29, 30].

To facilitate ease of use we make Exogene available as a Docker container which can be downloaded and run immediately, without requiring users to install third-party software (other than Docker itself) or to obtain specific versions of other resources.

## Conclusion

Exogene is an efficient and sensitive workflow for detecting viral integrations in human WGS, WES, RNA-Seq, and targeted capture paired-end sequencing data. We demonstrated Exogene’s accuracy via comparisons with long read validation for 6 HCC tumor samples, and demonstrated its applicability to targeted capture data by applying it to 426 previously studied tumor/normal pairs. Exogene’s read filtering and breakpoint detection strategies improve upon our previous workflow, yielding high confidence integration site coordinates. Exogene is freely available at github.com/zstephens/exogene. Additionally, we have made Exogene available as a Docker container to facilitate ease of use.

## Supporting information

**S1 Exogene logic got selecting representative alignments for multi-mapped reads**.

**S2 All HBV integrations in targeted capture samples**.

## Acknowledgments

We would like to thank the Mayo Clinic Center For Individualized Medicine.

## Footnotes

1 https://www.genome.jp/virushostdb/

2 www.hgsc.bcm.edu/sites/default/files/documents/IlluminaBarcodedPaired-EndCaptureLibraryPreparation.pdf

3 www.pacb.com/wp-content/uploads/SMRTbell-Library-Preparation-for-High-Fidelity-Long-Read-Sequencing-Customer-Training.pdf

